# Tau directs intracellular trafficking by regulating the forces exerted by kinesin and dynein teams

**DOI:** 10.1101/173609

**Authors:** Abdullah R. Chaudhary, Florian Berger, Christopher L. Berger, Adam G. Hendricks

## Abstract

Organelles, proteins, and mRNA are transported bidirectionally along microtubules by plus-end directed kinesin and minus-end directed dynein motors. Microtubules are decorated by microtubule-associated proteins (MAPs) that organize the cytoskeleton, regulate microtubule dynamics and modulate the interaction between motor proteins and microtubules to direct intracellular transport. Tau is a neuronal MAP that stabilizes axonal microtubules and crosslinks them into bundles. Dysregulation of tau leads to a range of neurodegenerative diseases known as tauopathies including Alzheimer’s disease (AD). Tau reduces the processivity of kinesin and dynein by acting as an obstacle on the microtubule. Single-molecule assays indicate that kinesin-1 is more strongly inhibited than kinesin-2 or dynein, suggesting tau might act to spatially modulate the activity of specific motors. To investigate the role of tau in regulating bidirectional transport, we isolated phagosomes driven by kinesin-1, kinesin-2, and dynein and reconstituted their motility along microtubules. We find that tau biases bidirectional motility towards the microtubule minus-end in a dose-dependent manner. Optical trapping measurements show that tau increases the magnitude and frequency of forces exerted by dynein through inhibiting opposing kinesin motors. Mathematical modeling indicates that tau controls the directional bias of intracellular cargoes through differentially tuning the processivity of kinesin-1, kinesin-2, and dynein. Taken together, these results demonstrate that tau modulates motility in a motor-specific manner to direct intracellular transport, and suggests that dysregulation of tau might contribute to neurodegeneration by disrupting the balance of plus- and minus-end directed transport.

**Synopsis and Graphical Table of Contents:** We isolated endogenous cargoes, along with a complement of kinesin-1, kinesin-2, and dynein motors, and reconstituted their bidirectional motility in vitro. We find that tau, a microtubule-associated protein that stabilizes microtubules in neuronal axons, directs bidirectional cargoes towards the microtubule minus end by tuning the balance of forces exerted by kinesin and dynein teams. These results suggest a general mechanism for regulating the transport of intracellular cargoes through modulating the relative activity of opposing motor teams.

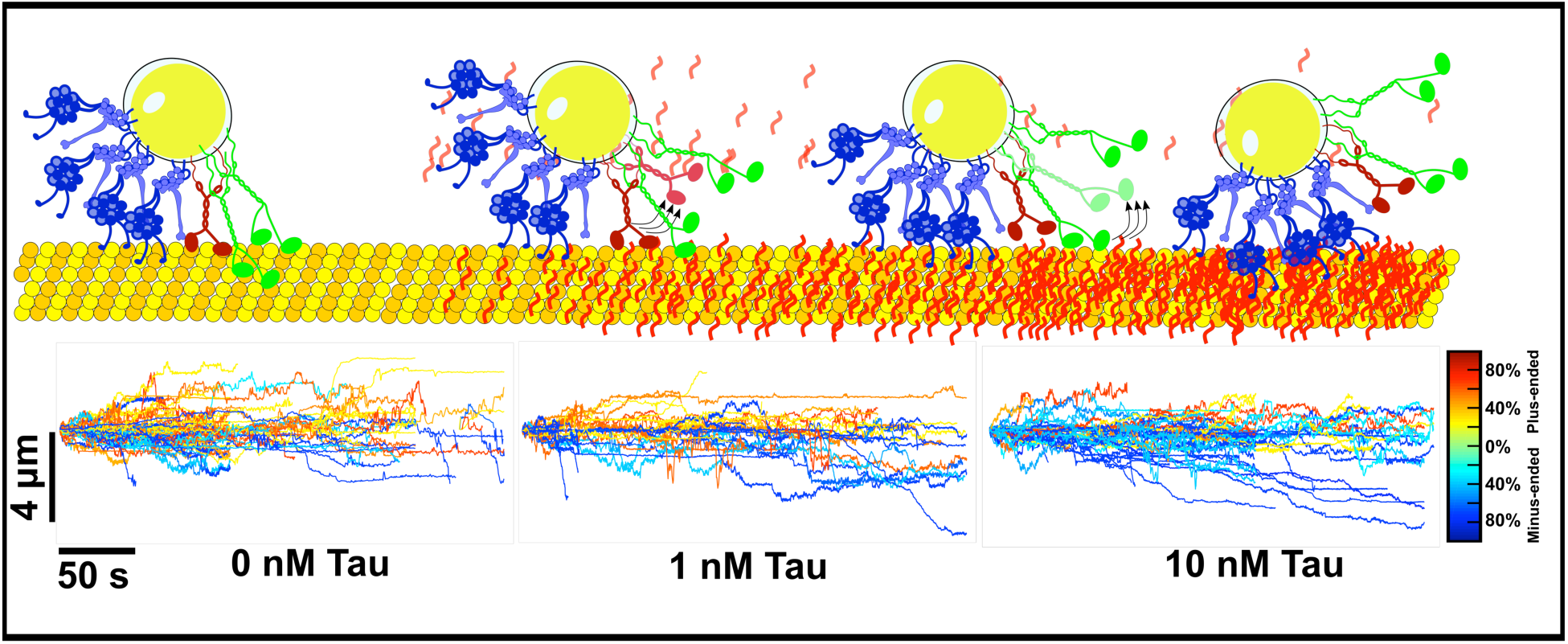

## Introduction

The motor proteins kinesin and dynein transport organelles, proteins, and mRNA along microtubules. Kinesin motors move towards the cell periphery and dynein moves towards the cell center. Many intracellular cargoes associate with both kinesin and dynein motors to enable movement in both directions along microtubules and to navigate around obstacles through the dense cellular environment**^1–3^**. Further, regulatory factors modulate the activity of kinesin and dynein to target cargoes to specific locations in the cell**^4, 5^**. Proper regulation of intracellular transport is required to maintain biosynthetic, degradative, and signaling pathways, and defects in trafficking are associated with developmental and neurodegenerative diseases**^6, 7^**.

One potential mechanism for regulating intracellular transport is through microtubule-associated proteins (MAPs) that bind along microtubules and modulate the interaction of motors with the microtubule surface. Tau is a neuronal MAP that bundles and stabilizes microtubules in the axon**^8, 9^**. Tau inhibits kinesin-1 motility in vitro by acting as an obstacle on the microtubule surface**^10, 11^**. In contrast, kinesin-2 and dynein are better able to navigate in the presence of obstacles and are only inhibited at high levels of tau**^10, 12, 13^**. By exerting differential effects on the motility of kinesin and dynein, tau might act to tune the balance of plus- and minus-end directed transport.

In support of a central role in regulating axonal transport, overexpression of tau in epithelial and neuronal cells results in mislocalization of mitochondria and other vesicular cargoes to the cell center by preferentially inhibiting plus-end directed movement**^12, 14–16^**. Tau knock-out mice exhibit age-dependent loss of neurons and defects in microtubule organization and axonal trafficking**^9, 17^**. In humans, tau mutations result in neurodegenerative disease**^18^** and tau dysregulation is implicated as a key driver of Parkinson’s**^17^** and Alzheimer’s**^19^** diseases.

Tau dynamics are regulated through post-translational modifications and isoform expression. Six tau isoforms are expressed in humans, varying in the number of microtubule-binding repeats and the length of the projection domain**^18^**. The length of the projection domain modulates tau’s effect on motor protein motility, with the shortest isoform exhibiting the strongest inhibition**^10^**. Over 85 putative phosphorylation sites have been identified for tau**^18^**, and tau phosphorylation modulates the dynamic equilibrium between static and transient binding to the microtubule**^20, 21^**. Tau phosphorylation is tightly regulated during development**^22^** and aberrant phosphorylation is linked to disease**^17, 23, 24^**.

To examine the influence of tau on organelle transport in the absence of cytosolic components, phosphorylation, or signaling events present in the cell, we isolated intracellular cargoes, along with their native transport machinery, and reconstituted their bidirectional motility along microtubules**^25, 26^**. Latex-bead-containing phagosomes are a useful model system for studying bidirectional transport. Their uniform size and density enables facile, high-purity isolation**^27^** and **^25, 28^**. The motor proteins and adaptors that transport phagosomes have been extensively characterized through proteomic and biophysical assays**^25, 27–30^**. Late phagosomes, as studied here, are transported by the microtubule motors kinesin-1 (KIF5B), kinesin-2 (KIF3A/B), and dynein-dynactin**^25, 29, 30^** and are positive for late endosomal markers such as LAMP1, Rab5, Rab7 **^27^** and motor adaptors including dynactin and Lis1**^25^** and KAP3**^31^.**

## Results

### Phagosomes are driven by teams of kinesin and dynein motors

To characterize the type and number of microtubule motors associated with isolated phagosomes, we used immunofluorescence and quantitative photobleaching. We performed three-color immunofluorescence on isolated phagosomes and found that 47% of phagosomes were positive for kinesin-1, kinesin-2, and dynein; while 17% were positive for kinesin-2 and dynein and 11% were positive for kinesin-1 and dynein. For a small fraction of phagosomes, we detected a single motor type **(Fig. 1b-d)**. Taking into account the likelihood that in some cases motors are present but not detected, these results indicate that most late phagosomes are transported by kinesin-1, kinesin-2, and dynein, in agreement with previous studies**^25, 29^**. To estimate the number of each type of motor bound to the phagosomes, we used stepwise photobleaching. For these assays, we labelled each motor using Alexa 647 in separate flow chambers to ensure similar conditions for each motor. We identified steps in the intensity traces using a step detection algorithm based on a two-sample t-test**^32^**. Single kinesin-1 motors were imaged with identical conditions to estimate the number of fluorophores conjugated to each motor **(Fig. S1d)**. These results indicate that ∼1-3 kinesin-1, 2-6 kinesin-2, and 3-10 dynein motors are bound to each cargo on average **(Fig. 1e)**, similar to the estimates for late endosomes isolated from mouse brain**^2^**.

**Figure 1.**
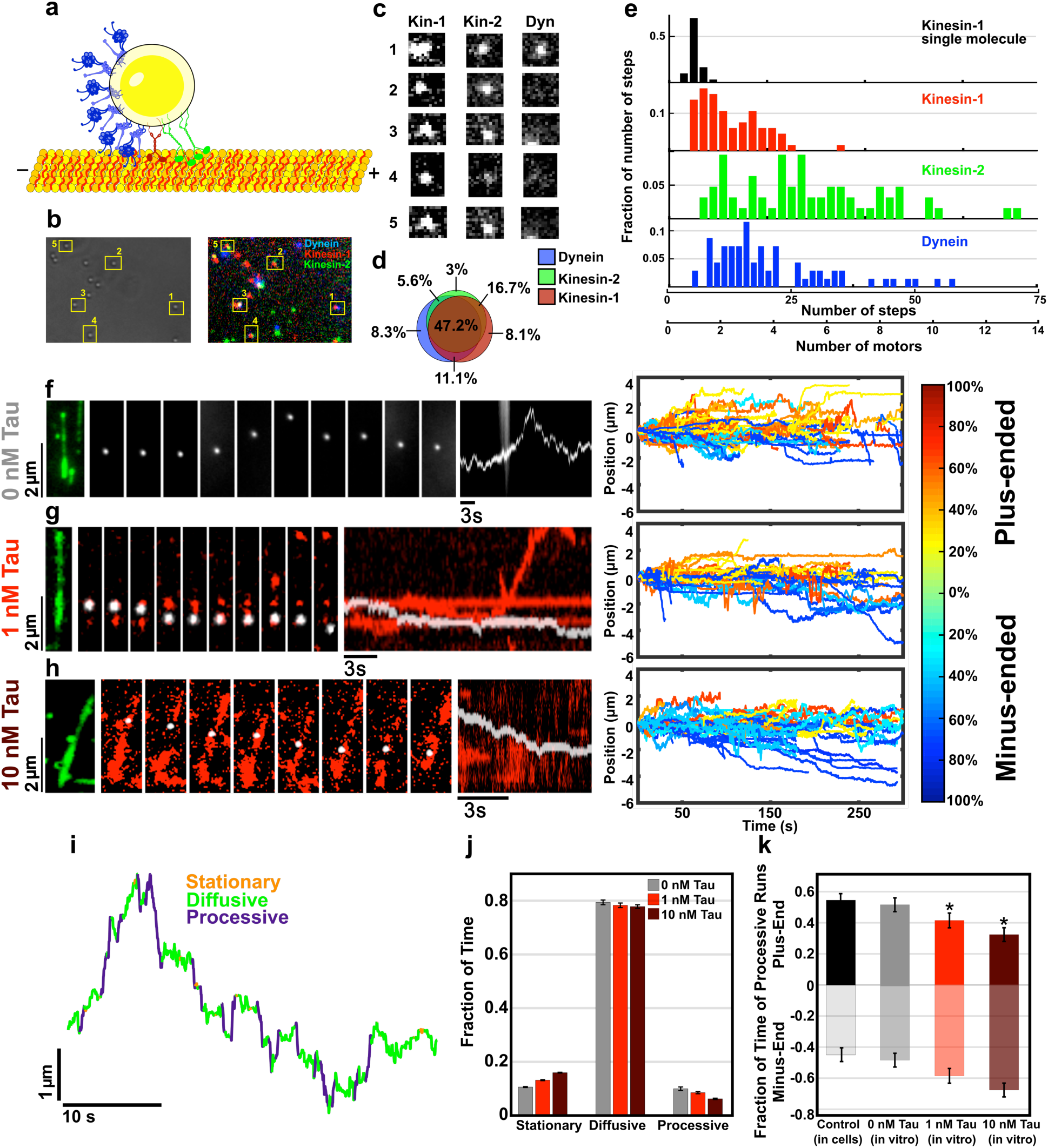
Tau directs the motility of isolated bidirectional cargoes towards the microtubule minus end. **(a)** Isolated phagosomes are transported by teams of kinesin-1 (red), kinesin-2 (green), and dynein (blue) motors. The microtubule-associated protein tau (red) binds along microtubules (yellow) and alters microtubule stability and motor-based transport. **(b and c)** Isolated phagosomes were probed for kinesin-1 (red), kinesin-2 (green) and dynein (blue) antibodies to identify the types of motors associated with each phagosome. Number of phagosomes (n=55). **(d)** Immunofluorescence indicates that approximately half of phagosomes are positive for kinesin-1, kinesin-2, and dynein. For other phagosomes, a subset of these motors was detected. **(e)** Stepwise photobleaching was used to estimate the number of motors associated with phagosomes. To estimate the number of fluorophores bound to a single motor, single kinesin-1 motors (rkin430-GFP) were analyzed under similar conditions. Photobleaching indicates that ∼ 1-3 kinesin-1, 2-6 kinesin-2, and 3-10 dynein motors are associated with each phagosome on average. N = 66 (kinesin-1), 63 (kinesin-2), and 69 (dynein) phagosomes. The number of steps for kinesin-2 was multiplied by a factor of 2 as kinesin-2 heterodimers contain one epitope, while kinesin-1 and dynein homodimers contain two epitopes for their respective antibodies per molecule. Example traces of stepwise photobleaching events are shown in **Fig. S1a, S1b, and S1c**. **(f-h)** Phagosomes containing fluorescent beads (white) move bidirectionally along polarity marked microtubules (green) in the absence and presence of tau (red). The middle panels show time-lapse images of the phagosome and tau movement, and kymographs are shown on the right. Isolated phagosomes move bidirectionally along microtubules, similar to their motility in cells. At low concentrations, tau binds in discrete clusters that diffuse along the microtubule lattice **(f)**. At higher concentrations tau decorates much of the microtubule lattice **(g)**. In the absence of tau, phagosomes exhibit approximately equal fractions of plus-(blue) and minus-(red) end directed movements. The color of the trajectory indicates the fraction of time of processive movements directed towards the microtubule plus- or minus- end. Tau biases the net directionality of cargoes towards the minus end in a dose-dependent manner. Number of trajectories, 0 nM tau (n_traj_=88, n_exp_=4), 1 nM tau (n_traj_=70, n_exp_=6), 10 nM tau (n_traj_=75, n_exp_=6). **(i)** Phagosome trajectories include periods of stationary, diffusive, and processive motility, characterized by the displacement between directional reversals (stationary: L_R_ < 10 nm; diffusive 10 nm ≤ L_R_ ≤ 400 nm; processive L_R_ ≥ 400 nm). The threshold for stationary runs was determined from the tracking error, and mean-squared displacement (MSD) analysis was used to determine the threshold for processive runs **(Fig. S2f)**. **(j)** The fraction of processive runs decreases in the presence of tau. **(k)** Tau increases the fraction of minus-end directed motility (*p<0.05). Error bars represent SEM.

### Tau biases directional motility of phagosomes towards the microtubule minus-end

The effect of tau on the motility of kinesin and dynein has been characterized for individual motors and multiple motors of one type **^10, 11, 16, 20, 33^**. Here, we investigate the potential role for tau in directing the motility of bidirectional cargoes driven by teams of both kinesin and dynein motors. Similar to their motility in living cells, isolated phagosomes move bidirectionally along microtubules, with periods of stationary, diffusive, and processive motility**^25^ (Fig. 1, Fig. S2)**. Using TIRF microscopy and subpixel resolution tracking **^34, 35^**, phagosomes were imaged with high spatiotemporal resolution (∼5 nm, 80 ms; and **SI Appendix, Fig. S2c)**. In the absence of tau, phagosomes exhibit approximately equal fractions of movement towards the microtubule plus- and minus-ends **(Fig. 1 f,k)**.

We focused on the shortest isoform of tau, 3RS, as it has been shown to have the strongest effects on motility**^10^**. At low concentrations (1 nM), tau forms small clusters that exhibit both static and diffusive motility along the microtubule **^36, 37^**. At higher concentrations (10 nM), tau decorates the entire microtubule lattice **(Fig. 1g,h)**. Tau alters the net directionality of transport by shifting motility towards the microtubule minus end. Diffusive movements are not affected by tau (Fig. S2). However, tau biases the processive movements of phagosomes towards the microtubule minus end, where 49% of processive motility is minus-end directed in the absence of tau compared to 59% (1 nM) and 65% (10 nM) when tau is present **(Fig. 1),** indicating that the density of tau on the microtubule surface determines the relative activity of kinesin and dynein.

### Tau preferentially inhibits kinesin to enhance dynein processivity on bidirectional cargoes

To examine how tau modulates motor processivity, we analyzed the length of individual runs between each reversal in the trajectories. Previous in vitro studies show that tau acts as an obstacle to decrease motor processivity, and that kinesin is inhibited more strongly than dynein by tau. When applied to bidirectional cargoes, these results suggest the hypothesis that tau will decrease kinesin run lengths sharply while dynein run lengths will also decrease but to a lesser extent. Instead, we observe that tau does not significantly alter the mean run length of plus-end directed runs, while the frequency of long processive runs towards the minus end increases in the presence of tau **(Fig. 2a).** This mechanism of regulation is specific to cargoes driven by both kinesin and dynein motors – tau reduced both the plus- and minus-end directed run lengths of bidirectional mRNA complexes driven by dynein alone**^16^**, and reduced the unidirectional run lengths of beads transported by kinesin**^11^** or dynein**^13^** alone. The length of individual processive runs is ∼200 nm **(Fig. 2)**, compared to several microns for cargoes transported by multiple kinesins or dynein, indicating that run lengths are governed by the interaction between opposing motors. Together these results suggest that for bidirectional cargoes driven by opposing motor teams, tau enhances dynein processivity by reducing the activity of opposing kinesin motors.

**Figure 2.**
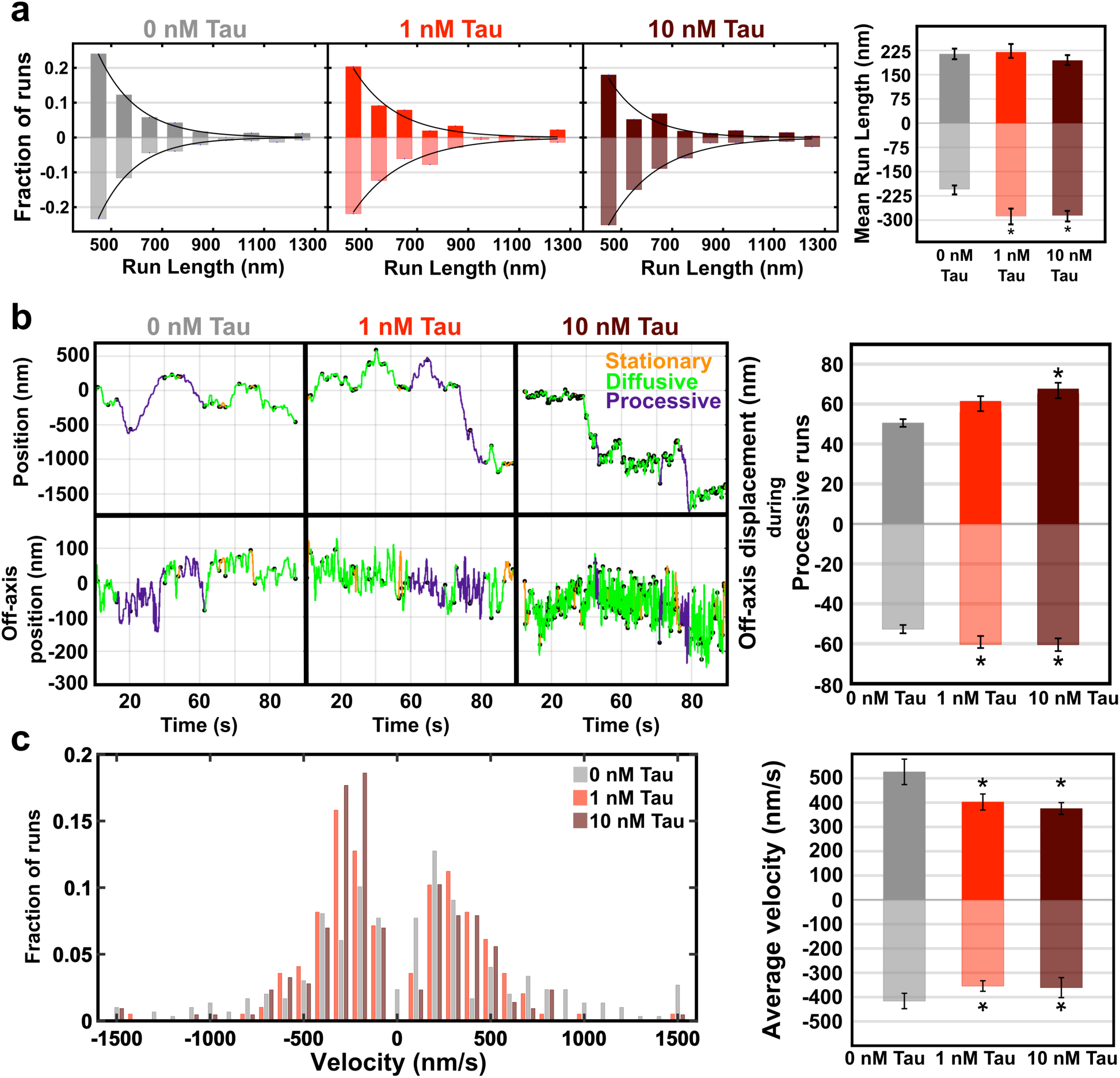
Tau increases the frequency of long dynein-directed runs. (**a**) Plus-end directed run lengths are largely unaffected by tau, while long processive runs towards the minus end are more frequent (*p<0.05). Number of processive runs, 0 nM tau (n=298), 1 nM tau (n=207), 10 nM tau (n=215). **(b)** Kinesin and dynein navigate around tau obstacles as indicated by increased off-axis displacements (*p<0.05). **(c)** Average velocities of phagosomes towards the microtubule plus and minus end are decreased in the presence of tau, accompanied by more frequent off-axis displacements (*p<0.05). Error bars represent SEM.

We next analyzed the off-axis displacements to investigate how teams of motors navigate on tau-decorated microtubules. Off-axis displacements increase in the presence of tau for both plus- and minus-end directed runs, indicating that both kinesin and dynein teams switch protofilaments to avoid obstacles on the microtubule **(Fig. 2b)**. Individual dynein and kinesin-2 motors can step to adjacent protofilaments**^33, 38, 39^**. For cargoes driven by multiple motors, sideways steps could also result from the binding and unbinding of motors to multiple protofilaments. Average velocities of phagosomes along the microtubule axis decrease with increasing tau concentration in both directions, consistent with tau acting as an obstacle to transport **(Fig. 2c)**.

### Dynein teams exert higher forces on tau-decorated microtubules

To test the hypothesis that tau reduces the forces exerted by kinesin motors to enhance dynein motility, we measured forces exerted by phagosomes using an optical trap. Teams of dynein and kinesin motors exert forces up to 9 pN, with an approximately equal fraction of plus- and minus-end directed events consistent with their motility **(Fig. 1f, 3a, S3)**. The plus-end directed forces show a peak ∼4 pN similar the unitary stall force for kinesin-1 and kinesin-2 **(Fig. 3a, S3c)**. Lower force events (∼1.4 pN and 2.3 pN) may be due to detachments before the motors reached stall or opposing loads from dynein**^40^**. Dynein forces exhibit peaks at ∼1 pN intervals consistent with events driven by multiple dynein motors, each exerting ∼1 pN **^25, 28^**. Tau decreases the frequency and magnitude of kinesin-driven forces, indicating that both fewer kinesins are able to engage **^11^** and that the engaged kinesins exert less force in the presence of tau **(Fig. 3b, S3d)**. Correspondingly, minus-end directed force events are more frequent and shifted towards higher magnitudes, suggesting that dynein teams are able to exert higher forces when force generation by opposing kinesin motors is inhibited by tau.

**Figure 3.**
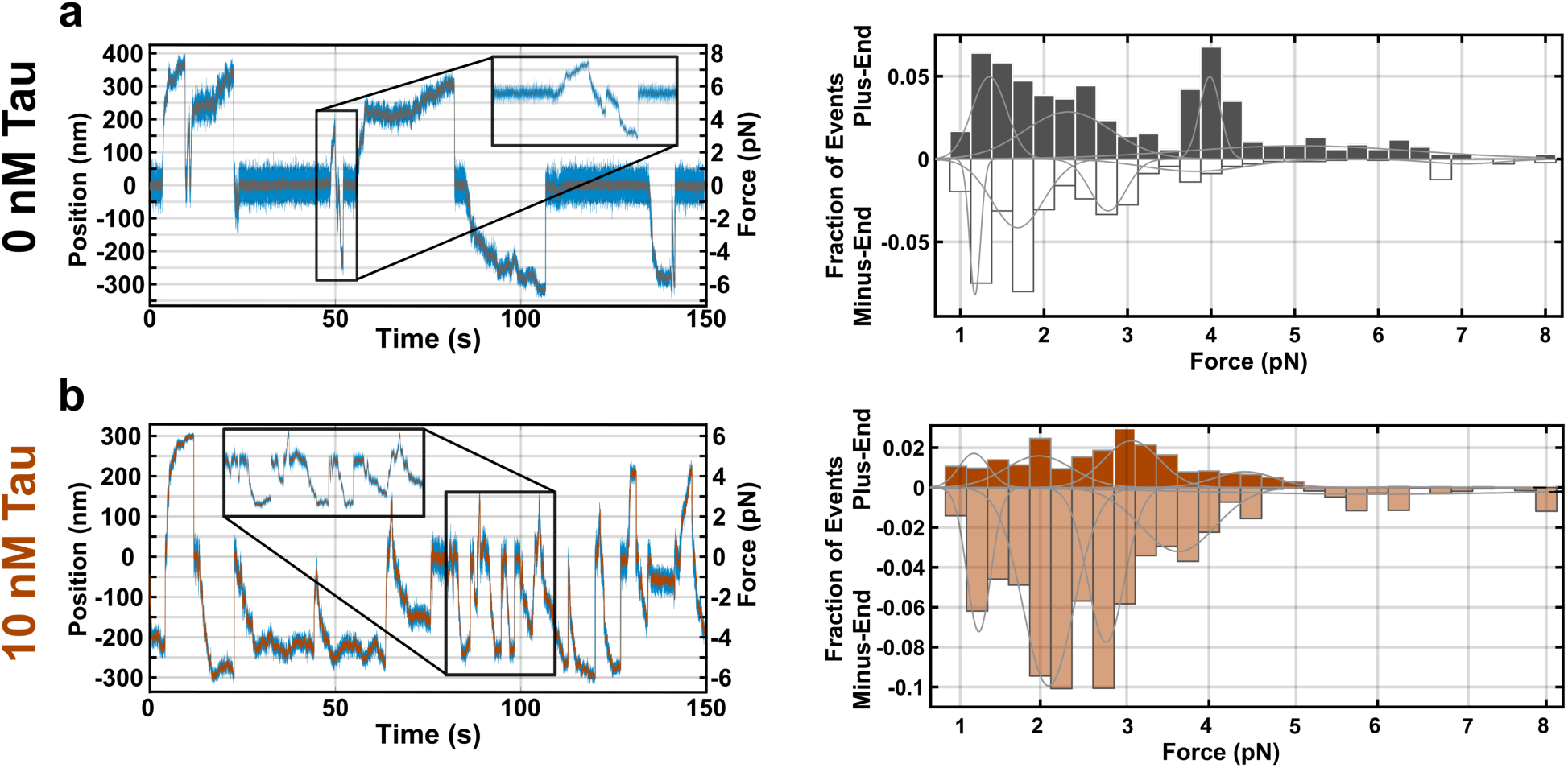
Dynein motors exert higher forces in the presence of tau. **(a)** Isolated phagosomes were positioned on polarity-marked microtubules and the forces were measured using an optical trap. Force traces were acquired at 2 kHz and median filtered to 20 Hz. The histograms include the maximum forces for all trap displacements greater than 300 ms in duration (n=949 events from 60 recordings, 7 independent experiments). Plus-end directed force traces are consistent with stall events due to single kinesin-1 and -2 motors ∼4-5 pN, as well as events where motors detach before reaching their maximal force and rare events driven by multiple kinesins. Minus-end directed forces indicate events driven by several dynein motors, each exerting ∼1.2 pN. The Bayesian Information Criterion was used to determine the optimal number of components to describe the force histograms **(Fig. S3g,h)**. Mean forces of the multicomponent fits for plus-end directed forces are 1.36 pN, 2.3 pN, 4.12 pN, 5.14 pN, and for minus-end directed forces are 1.17 pN, 1.75 pN, 2.75 pN, 3.81 pN, 7.04 pN. **(b)** In the presence of 10 nM tau, the frequency of force events in the plus-end direction are greatly reduced (n=1913 events from 81 recordings, 10 independent experiments). Kinesin exerts less force, and events greater than 5 pN are rare. In response, dynein exerts higher forces and dynein-directed forces are more frequent. Mean forces of the multicomponent fits for plus-end directed forces are 1.2 pN, 1.99 pN, 3.07 pN, 4.43 pN and for minus-end directed forces are 1.27 pN, 2.1 pN, 2.8 pN, 3.66 pN, 6.03 pN.

### Tau regulates transport through motor-specific tuning of processivity

Previously, a mathematical model demonstrated that a stochastic tug of war between teams of opposing motor proteins in the absence of external regulation results in bidirectional motility similar to the trajectories we observe**^41^**. We extended this model to simulate the motility of cargoes transported by kinesin-1, kinesin-2, and dynein **(SI Appendix)**. Optical trapping measurements show that maximum forces exerted by teams of plus-ended motors is ∼9 pN indicating that up to 3 kinesin motors are engaged at any one time. Forces of up to 9 pN are exerted in the minus-end direction, indicating that ∼ 10 dynein motors can be engaged simultaneously, assuming each dynein exerts ∼1 pN **(Fig. 3a,b)**. These measurements indicate that a subset of the total number of motors bound to the cargo are engaged at any one time **(Fig. 1b-e)**. By modeling the motility of cargoes transported by one kinesin-1, two kinesin-2, and ten dynein motors, we found similar trajectories to those observed for isolated phagosomes **(Fig. 4, and Table S1)**. We estimated the effect of tau on the unbinding rates of kinesin-1, kinesin-2, and dynein using data from in vitro experiments**^41, 42^ (Fig. 4e, and SI Appendix, Fig. S4a)**. Consistent with our experimental results, the number of minus-ended trajectories increases with increasing tau concentration **(Fig. 4a, d)**. Further, the model results agree with the observation that the plus-end directed run lengths are not largely affected, but minus-end directed run lengths increase due to tau **(Fig. 4b, c)**. For comparison, we modeled the effect of tau on cargoes where only kinesin-1 or kinesin-2 is active. When both kinesin-1 and kinesin-2 are engaged, the effect of tau on the unbinding rate of kinesin-1, kinesin-2, and dynein closely follows the results from single molecule experiments**^10, 20, 33^ (Fig. S4a)**. In simulations of cargoes driven by kinesin-1 and dynein, the unbinding rate of kinesin-1 was predicted to be less sensitive than observed for single molecules**^10^** to obtain similar trajectories. For cargoes driven by kinesin-2 and dynein, the unbinding rates of kinesin-2 were predicted to be much more sensitive to tau than observed for single molecules **(SI Appendix, Fig. S4, Table S2)**, indicating that the model where both kinesin-1 and kinesin-2 contribute to phagosome motility is most consistent with our results. In agreement, most phagosomes co-purify with both kinesin-1 and -2 (Fig. 1), and inhibiting either kinesin-1 or kinesin-2 reduces the motility of phagosomes in cells**^25^**. These results suggest that recruiting kinesin-1 to a cargo may be a way to confer sensitivity to regulation by tau.

**Figure 4.**
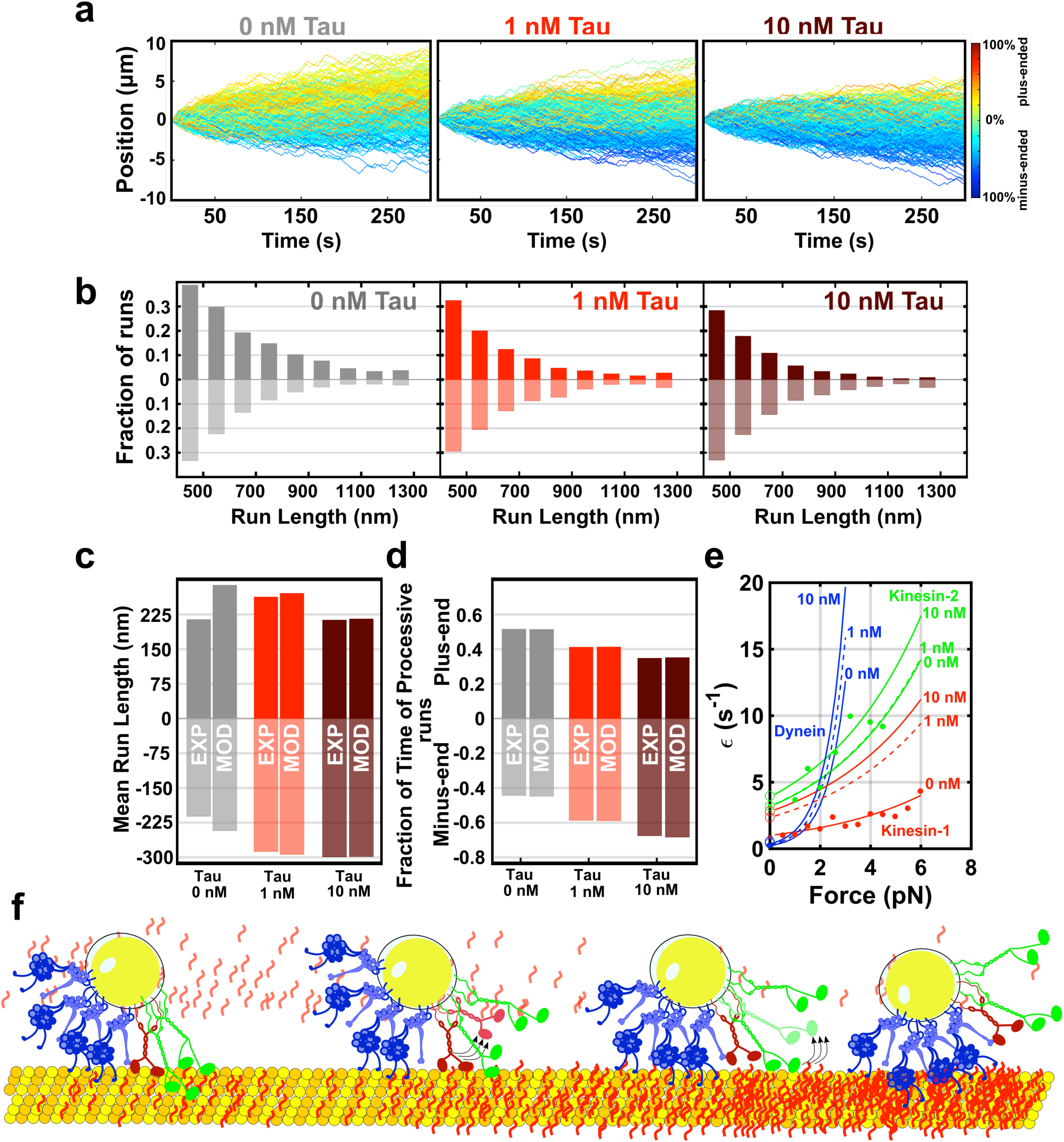
Mathematical modeling indicates that tau biases transport towards the minus end by preferentially inhibiting kinesin-1 processivity. The model of bidirectional motility developed by Müller, Klumpp, and Lipowsky **^41^** was modified to describe the interaction of kinesin-1, kinesin-2, and dynein motors. Single motor parameters such as stall force, detachment force, unbinding and binding rates were determined from single molecule data **^41, 42^.** The effect of tau was modeled by increasing the dissociation rate of kinesin-1, kinesin-2, and dynein **(^10, 20^; Table S1,2).** Circles represent the unbinding rates determined from single-molecule experiments**^10, 20, 33^**. Trajectories were simulated for cargoes transported by one kinesin-1, two kinesin-2, and ten dynein motors. **(a)** Simulated trajectories show bidirectional motility similar to isolated phagosomes. Lines indicate trajectories with net directionality towards the microtubule plus (blue) and minus (red) end. As tau concentration increases, the fraction of minus-end directed motility increases. Number of trajectories, 0 nM tau (n=400), 1 nM tau (n=400), 10 nM tau (n=400). **(b-c)** Similar to the experimental trajectories, the run lengths of dynein-directed runs are increased by tau. **(d)** The model indicates that tau regulates the directionality of cargoes by inhibiting the processivity of kinesin-1 more strongly than kinesin-2 and dynein. **(e)** Data from single-molecule experiments was used to estimate the sensitivity of motor unbinding rates to load **^42, 57–60^** (closed circles) and tau **^10, 20, 33^** (open circles) **(Fig. S4a)**. Kinesin-1 is more sensitive than kinesin-2 and dynein to tau. **(f)** Through preferentially inhibiting kinesin-1, tau reduces the forces exerted by kinesin teams, resulting in enhanced dynein processivity and force generation. By targeting specific motors, tau’s effect on motility can be tuned to each cargo depending on the makeup of the motor teams driving its transport.

## Discussion

By reconstituting the bidirectional motility of endogenous organelles in the presence of MAPs, we show that tau tunes the balance of plus- and minus-end directed transport through motor-specific regulation of processivity. On bidirectional cargoes driven by both kinesin and dynein, preferential inhibition of kinesin results in fewer engaged kinesins such that the kinesin team exerts less force. Dynein processivity is enhanced in response to reduced opposing forces, shifting transport towards the microtubule minus end. Regulation of transport by tau suggests a general mechanism, where small perturbations in the force generation or processivity of opposing motors results in biasing the net direction of transport. Interestingly, inhibition of either kinesin or dynein via function-blocking antibodies, dominant-negative constructs, or RNAi often halts motility in both directions**^25, 43, 44^**. This suggests that regulation of transport may instead rely on subtle changes to motor activity as observed here.

Intracellular cargoes are often transported by dynein and multiple kinesin members. Kinesin-1, kinesin-2, and dynein interact to transport late endosomes**^2, 29^**. Kinesin-1, kinesin-3, and dynein associate with mitochondria and synaptic vesicle precursors**^45, 46^**. Employing multiple kinesin members on a single cargo might be a way to target tau regulation to specific cargoes. Mathematical modeling suggests that for cargoes transported by kinesin-1, kinesin-2, and dynein, regulation by tau is primarily mediated through kinesin-1 due to its high sensitivity to tau **(Fig. 4 and S4)**. These results imply that cargoes transported primarily by kinesin-1 and dynein, such as early endosomes, might be more strongly regulated by tau than cargoes where kinesin-2 is also present like late endosomes and phagosomes.

Further, the interaction between tau, motors, and microtubules is regulated through tau isoform expression and post-translational modifications. Six tau isoforms, generated through alternative splicing, differ in the number of microtubule-binding repeats and length of the acidic projection domain that extends from the microtubule surface**^18, 47^**. The shortest isoform of tau inhibits kinesin and dynein motility more strongly than longer isoforms**^10, 11, 48^**. Tau is modified by a variety of post-translational modifications, and over 85 putative phosphorylation sites have been identified **^49–51^**. While the function of most of these sites is unknown, phosphorylation of tyrosine 18 weakens tau’s affinity to microtubules and in turn the inhibition of kinesin-1 motility**^17, 21^**. Thus, multiple layers of regulation allow the cell to tune the local density and dynamics of tau to control trafficking in specific subcellular regions**^15, 52^**.

Taken together, these studies suggest that in addition to its role in organizing the microtubule cytoskeleton**^53, 54^**, tau has a central role in controlling the motility of motor proteins to direct trafficking. Further, different motors have varying sensitivity to tau, enabling tau regulation to be targeted to specific cargoes. Our observations indicate that defects in tau expression, localization, and binding dynamics would significantly alter the balance of plus- and minus-end directed transport, contributing to tau’s role in neurodegenerative diseases**^55^**.

## Methods

### Cell culture and phagosome purification

J774A.1 cells (ATCC) were plated in four 100 mm cell culture dishes and cells were grown in a 37^o^C incubator at 5% CO_2_ to ∼ 80% confluency in DMEM media (Thermo Fisher Scientific), supplemented with 10% FBS (Thermo Fisher Scientific) and 1% Glutamax (Thermo Fisher Scientific). Fluorescent latex beads (200 nm diameter, blue fluorescent, ThermoFisher Scientific) were coated with BSA and incubated with J774A.1 mouse macrophage cells. Larger beads (500 nm diameter, yellow-green fluorescent, ThermoFisher Scientific) were used for optical trapping to enable adequate forces to be exerted. Phagosomes were isolated as described **^26^.** After a 90-min chase, cells were washed with PBS, collected by a cell scraper and washed again with cold PBS, where after each wash the cells were centrifuged at 2000 RPM in 4^°^C for 5 min and resuspended in PBS. After the second wash, cells were resuspended in motility assay buffer (MAB; 10 mM Pipes, 50 mM K-acetate, 4 mM MgCl_2_, 1 mM EGTA, pH 7.0) supplemented with protease inhibitors, 1 mM DTT, 1 mM ATP, and 8.5% (w/v) sucrose solution in MAB and lysed with a Dounce homogenizer. Following the lysis step, cell lysate was centrifuged at 2000 RPM in 4^°^C for 10 min and the supernatant (homogenate) was mixed with an equal volume of 62% sucrose. Sucrose solutions were supplemented with ATP and protease inhibitors and loaded in ultracentrifuge tubes (16 x 102 mm – Beckman no. 344061) in the following order: 62% (3.61 mL), homogenate (∼ 1 mL), 35 % (2.91 mL), 25% for 500 nm beads or 30 % for 200 nm nm beads (2.91 mL), and 10 % sucrose (2.91 mL). The sucrose gradient was centrifuged in a swinging bucket rotor (SW32.1 – Beckman) at 24000 RPM for 72 minutes. After the spin, the fraction containing phagosomes appears as a thin band at the interface between 25% and 30% sucrose (for 200 nm beads) and between 25% and 10% sucrose (for 500 nm beads).

### Tau expression and purification

Wild-type 3RS-tau was purified as described^48^. Briefly, wild-type 3RS tau was expressed in BL21-CodonPlus DE3)-RP E. coli cells (Strategene) using pET vector system (Novagen). The bacterial pellet was ground under liquid nitrogen, then boiled and clarified via centrifugation. The supernatant was passed through a 0.22 μm filter, and purified by consecutive Q Sepherose and SP Sepherose Fast Flow columns (Sigma). Purified tau was dialyzed in BRB80 at room temperature and its purity was assessed by SDS-PAGE **(Fig S5)**. Tau concentration was determined by BCA assay (Pierce) using desalted, lyophilized 3RS-Tau as standards.

### Polarity-marked microtubules

Bright microtubule seeds were prepared by mixing 25% alexa 647 labeled tubulin and 75% unlabeled tubulin in BRB80 (80 mM K-PIPES, 1 mm MgCl_2_, 1mM EGTA, pH 6.8) to a final concentration of 5 mg/mL supplemented with 1 mM GTP (Sigma Aldrich) and polymerized at 37^°^C for 20 min. After polymerization, bright microtubule seeds were dissolved in cold BRB80 supplemented with 1 mM GMPCPP (Jena Bioscience) and 2 mM MgCl_2_. This mixture was incubated at 37^°^C for 30 min and then pelleted at 100,000 x g for 10 min at 4^°^C in TLA100 rotor (Beckman). The pellet was resuspended in 70 uL cold BRB80 and incubated on ice for 20 minutes for depolymerization. The solution was centrifuged at 100,000 x g for 10 min at 4^°^C and resuspended with 2 mM MgCl_2_ and 1 mM GMPCPP. The pellet was then resuspended in BRB80 and aliquoted in 5 μL stocks and flash frozen. To prepare polarity marked microtubules, GMPCPP seeds were warmed at 37^°^C for 1 min to which 8% labeled tubulin was added in 92% unlabeled tubulin in BRB80 and 1 mM GTP to a final concentration of 2 mg/mL. After 25 min incubation, these microtubules were stabilized with 20 μM taxol (Cytoskeleton) and incubated for another 25 min at 37^o^C.

### In vitro motility assay

Silanized coverslips were mounted on glass slides using vacuum grease and double sided tape to form 20-25 μL flow chambers. After flowing 1 chamber volume of anti-β-tubulin (2.5:50 in BRB80), the chamber was surface treated with F-127 (Sigma) to block any non-specific binding. Diluted microtubules (2:50 in BRB80 supplemented with Taxol) were flown through the chamber, following a wash with two chamber volumes of BRB80 supplemented with 20 μM taxol. Purified phagosomes were added to the chamber, supplemented with 0.2 mg/mL BSA, 10 mM DTT, 1 mm MgATP, 20 μM taxol, 15 mg/mL glucose, ≥2000 units/g glucose oxidase, ≥6 units/g catalase, and 1 mg/mL casein. For tau experiments, microtubules were mixed with Alexa 568 labeled tau and incubated in the flow chamber for 30 minutes. Phagosomes, supplemented as above and additionally with tau (1 nM or 10 nM) were added to the chamber and imaged using Total Internal Reflection Fluorescence (TIRF) microscopy at 80 frames per second.

### Subpixel tracking and analysis

Motility was analyzed using FIESTA (Fluorescence Image Evaluation Software for Tracking and Analysis) **^35^**. Position vs time trajectories from FIESTA tracking were compared to kymographs to validate the automated tracking. Stationary phagosomes were tracked to estimate the localization uncertainty **(Figure S1)**. A Savitsky-Golay filter (window size of 1.2 s with a 1^st^ degree polynomial) was applied to the trajectory, and runs were identified as the displacement between two reversals in a trajectory. Runs were categorized as stationary (0 nm > |L_rev_| ≥ 10 nm) corresponding to the tracking uncertainty of ∼ 6 nm, diffusive (10 nm ≥ |L_rev_| > 400 nm) or processive (|L_rev_| ≥ 400 nm). Mean-squared displacement analysis was used to determine the threshold for processive runs **(Figure S1E)**. The 1-dimensional mean squared displacement (MSD = 2Dt^α^) was calculated using internal averaging, where D is the diffusion coefficient, t is the time interval, and α is the scaling exponent corresponding to stationary (α=0), diffusive (α=1), and processive (α=2) movement **^56^**. To validate the thresholds for stationary, diffusive and processive motility, we checked that the fraction of plus- and minus-end directed motility was approximately equal for runs characterized as diffusive: 0 nM tau (49.6 % plus-ended and 50.4% minus-ended), 1 nM tau (50.1% plus-ended and 49.9% minus-ended) and 10 nM tau (47.9% plus-ended and 52.1% minus-ended) **(Figure S1F)**.

### Mathematical Modeling

Müller, Klumpp, and Lipowsky developed a model to describe vesicle transport driven by two teams of opposing motor proteins **^41^**. Directional switches are a result of the stochastic binding and unbinding kinetics of the motors in the absence of external regulation. We extended the MKL model to describe transport by teams of kinesin-1, kinesin-2, and dynein motors **(SI Appendix)**.

### Optical trapping

The optical trap was built on an inverted microscope (Eclipse Ti-E: Nikon) with a 1.49 NA oil-immersion objective. The beam from a 10W, 1064 nm laser (IPG Photonics) was expanded to overfill the back aperture of the objective. A position-sensitive detector (Thorlabs), positioned conjugate to the back focal plane of the condenser, provided a measurement of the bead displacement and force. The optical trap stiffness and positional calibration was determined by fitting a Lorenzian function to the power spectrum of bead thermal fluctuations. Motility assays were performed as described above, with the exception that 500-nm beads were used to allow adequate forces to be exerted by the optical trap.

### Immunofluorescence of motor proteins on isolated phagosomes

Phagosomes were added to silanized coverslips and incubated for 3 hours. Following the incubation step, the chamber was surface treated with F-127 to reduce any non-specific binding. Primary mouse antibody against kinesin-1 (MAB1614, EMD millipore) and primary rabbit antibody against dynein (sc-9115, Santa Cruz) were added to the flow chambers and incubated for 1 hour at room temperature. After washing the flow chamber twice (with 1x BRB80), secondary antibodies against mouse (Alexa 647 - A21235, Thermo Fisher Scientific) and rabbit (Atto 488 - 18772, Sigma-Aldrich) were incubated with the sample for 45 minutes. The flow chamber was then washed (x3) with 1x BRB80. Kinesin-2 primary antibody conjugated to Alexa 568 (Abcam, ab24626 and Thermo Fisher Scientific, A20184) was added to the chamber and incubated for 2 hours in the dark. The sample was then washed 3 times with 1x BRB80 before imaging. The phagosomes were imaged in brightfield and epifluorescence to determine if signal by the fluorophore was from the phagosomes.

### Stepwise photobleaching

Three separate silanized coverslips were incubated with phagosomes and surface treated with F-127. The coverslips were incubated with primary antibodies to kinesin-1 (MAB1614, EMD millipore), kinesin-2 (Abcam, ab24626), and dynein (MAB1618, EMD millipore) respectively for 2 hours at room temperature, washed three times with BRB80. The coverslips were incubated with Alexa 647 secondary antibodies for 45 minutes at room temperature, washed three times in BRB80, and imaged.

To perform stepwise photobleaching on kinesin-1 (rkin430-GFP), protocol similar to the in vitro motility assay was performed. Prior to adding the kinesin-1 on the microtubules, it was incubated with a kinesin-1 primary (MAB1614, EMD millipore) and Alexa 647 secondary antibodies (A21235, Thermofisher). Kinesin-1 was bound to the microtubules by using AMPNP and washed two times with TBRB80, and imaged using TIRF microscopy.

Imaging was performed using a 647 laser (3 mW) with an exposure time of 500 msec. Only steps greater than 30 (a.u.) were considered. The unitary step size - the size of the step due to the phobleaching of a single molecule - was estimated as the mean step size for all the steps detected for kinesin-1, kinesin-2, and dynein. The number of steps was calculated by dividing the difference between the initial and final intensity by the unitary step size**^32^ (Fig. S1).**

### Statistical analysis

All the data is presented as standard error of mean (SEM). The value of n and the number of independent experiments are mentioned in the figure legends. Statistical analysis was performed using MATLAB (Mathworks Inc.). Statistical analysis was performed using One-Factor ANOVA and student’s t-test (for off-axis displacement and average velocity), bootstrapping and Efron’s percentile method (for fraction of time of processive runs and fraction of time of force events), and Klomogrov-Smirnov test (for processive run lengths).

## Acknowledgements

The authors thank Dr. Gary Brouhard and Lynn Chrin for providing reagents and technical assistance. F.B. is a Feodor Lynen Fellow of the Alexander von Humboldt Foundation. This work was supported by NSERC (Discovery Grant) and FRQNT (New Investigator) grants to A.G.H. and NIH grant (R01 GM101066) to C.L.B.

## Author Contributions

A.R.C and A.G.H designed research. A.R.C performed the experiments. F.B. wrote the mathematical model. C.L.B contributed reagents. A.G.H and A.R.C analyzed the data, and wrote the manuscript. A.G.H conceptualized the project and supervised the work.

## Supplementary Information

Supplemental Information for this article includes five supplemental figures, four supplemental tables, and four movies.

